# Factors Influencing the Detection of Antibacterial Resistant *Escherichia coli* in Faecal Samples from Individual Cattle

**DOI:** 10.1101/2021.11.07.467618

**Authors:** Andrea Turner, Hannah Schubert, Emma F. Puddy, Jordan E. Sealey, Virginia C. Gould, Tristan A. Cogan, Matthew B. Avison, Kristen K. Reyher

## Abstract

**Aims:** To investigate whether on-farm antibacterial usage (ABU), environmental antibacterial resistant (ABR) *Escherichia coli* prevalence, and sampling and sample handling methodologies are associated with ABR *E. coli* positivity in individual faecal samples from dairy heifers.

**Methods and Results:** Three hundred and sixty-four heifers from 37 farms were sampled via rectal or faecal pat sampling. Samples were stored at -80°C for variable periods before microbiological analysis. Data analysis was through a multilevel, multivariable logistic regression approach.

Individual rectal samples had increased odds of positivity for amoxicillin, cefalexin and tetracycline-resistant *E. coli*. Sample storage for 6-12 month was associated with decreased odds of finding amoxicillin and tetracycline-resistant *E. coli*. On-farm ABU had little influence, and environmental ABR *E. coli* prevalence had no significant influence on on the odds of sample-level positivity for ABR *E. coli*.

**Conclusions:** Sampling methodology and sample handling have a greater association than on-farm factors with the detection of ABR *E. coli* in individual faecal samples from dairy heifers.

**Significance and Impact of the Study:** Sampling and storage methodologies should be considered carefully at the point of designing ABR surveillance studies in livestock and their environments and, where possible, standardised between and within future studies.

## Introduction

There is an increasing expectation on livestock industries to reduce the risk of antibacterial resistance (ABR) selection and so to reduce transmission of ABR bacteria from animals to humans. It is known that bacteria such as *Escherichia coli* can be transmitted between livestock and humans (Carattoli, 2008, Ho *et al*., 2010). This can occur in several ways including direct contact, through food or via transmission from the environment. To understand the dynamics of ABR transfer within animals, between animals and between animal and human populations, it is crucial to also understand which factors are responsible for the presence of ABR in animal populations. It is therefore essential to be able to measure ABR prevalence within animal populations and their environments using reliable and reproducible methods, and, for logistical reasons, to identify samples that are representative and yet relatively easy to obtain.

Animals of different ages may show differing ABR prevalence among colonising flora. Furthermore, ABR prevalence in cattle environments has been found to be higher in areas where youngstock are housed when compared to environments of the adult herd (Hancock *et al*., 1997, Horton *et al*., 2016, Schubert *et al*., 2021) and bacterial clones have been shown to transmit between groups of animals of different ages, even on farms with good internal biosecurity (Agren *et al*., 2018). These living conditions and transmission pathways are not easily separated, however; in most modern dairy systems youngstock are grazed on pastures also used for adult cattle and manure from youngstock housing contributes to the material spread on pasture. For these reasons it is important that faecal sampling methodologies for the detection of ABR prevalence are validated for dairy youngstock populations as well as the adult herd.

Numerous studies have been conducted investigating risk factors for the presence and prevalence of ABR in *E. coli* isolates on dairy farms (Snow *et al*., 2012, Watson *et al*., 2012, Brunton *et al*., 2014, Duse *et al*., 2015, Gonggrijp *et al*., 2016, Santman-Berends *et al*., 2017). Similarly, several studies have shown that sample handling and environmental factors, including ambient temperature at the time of sample collection (Schubert *et al*., 2021), influence the sensitivity of testing methods to detect bacteria in environmental and faecal samples from livestock (Oladeinde *et al*., 2014, Oliver *et al*., 2016, Schubert *et al*., 2021). It should also be considered that the detection of ABR in animal faecal samples could be affected by sample collection methodology, which varies between studies. Across a range of studies, faecal samples have been collected either directly from animals (Brunton *et al*., 2014, Pereira *et al*., 2014) or from faecal pats or faecally contaminated areas in the animals’ environment (Dunlop *et al*., 1999, Watson *et al*., 2012, Gonggrijp *et al*., 2016, Horton *et al*., 2016). In general, sampling from faecal pats offers a simple method for sample collection which can be done by lay people with minimal stress to animals, minimal health and safety concerns and without the need for ethical approval for animal handling. This method, however, often makes it difficult or impossible to associate samples with individual animals. In some studies, both direct and faecal pat sampling methods have been applied, occasionally interchangeably, when direct sampling of individual animals is unfeasible due to logistics or where there are safety concerns for the animal or researcher (Hancock *et al*., 1997). To the authors’ knowledge, only one study has attempted to validate the use of both faecal pat and individual animal sampling to determine ABR levels in animal populations, but the pat samples were from the pen floor and not directly linked with individual animals (Wagner *et al*., 2002).

To address some of these questions, this study was designed to assess how commonly used sampling methodologies and sample handling, alongside other potential drivers, affect the detection of ABR *E. coli* isolates in faeces from dairy heifers on 37 dairy farms in South West England. This study investigated the hypothesis that sampling methodologies and sample handling would affect ABR prevalence in these samples. Specifically, the effects of collecting faeces directly from the rectum as opposed to from faecal pats, timing of post-sample microbiological analysis, antibacterial usage (ABU) on the farm and ABR prevalence in environmental samples previously collected from the farm were analysed to assess their associations with ABR detection in individual heifer faecal samples.

## Materials and Methods

### Farm recruitment and ethical approval

Fifty-three dairy farms located in South West England had previously been enrolled into a wider study by the same research group: the OH-STAR study (Schubert *et al*., 2021). This was a convenience sample of farms recruited through personal contacts, local veterinary practices, and milk processors. These 53 farms were assessed for suitability to enrol in the parallel study reported here. All farmers gave fully informed consent to participate. Ethical approval was obtained from the University of Bristol’s Faculty of Health Sciences Research Ethics Committee (ref 41562).

### Sampling

Thirty-seven farms contributed to this study (the remaining 16 farms in the OH-STAR study either did not rear heifers or could not be sampled for this study).

As part of the OH-STAR study, farms were visited monthly from January 2017 until December 2018. On these visits, samples of faecally contaminated environments (environmental samples) were collected from several locations on each farm (collecting yard, fields and sheds) using sterile over-boot socks. Environmental samples (>4500 in total across the whole study) were then analysed for the presence of *E. coli* resistant to one or more of 5 commonly used antibacterials (cefalexin, amoxicillin, tetracycline, streptomycin and ciproflaxacin). The sampling methodology for these environmental samples is described more fully elsewhere (Schubert *et al*., 2021). Sample-level positivity data for 3 of the test antibacterials (cefalexin, amoxicillin and tetracycline) from the environmental samples collected as part of the OH-STAR study, were included in the modelling of the study this paper describes.

In addition, for the present study, 37 farms were visited on a single occasion on which approximately 10 (range 2-22) heifers from each farm had individual faecal samples collected at a single time point. The heifers selected for individual sampling were those closest to 18 months of age at the time of sampling and accessible to the researcher during the sampling period (age range 14-22 months, mean 18). These ‘individual heifer’ sampling visits happened between June and September 2018. Importantly, the wider OH-STAR study had collected environmental samples in areas close to these heifers monthly throughout their lives, as part of its wider environmental sample collection strategy.

Individual heifer samples were collected in 2 ways:

1. If, during the required time period, the selected heifers were being examined rectally by a veterinary surgeon for fertility or other health assessment purposes, the rectal glove used for the examination was passed immediately to the researcher by the veterinary surgeon. Faeces was immediately transferred from the rectal glove into a universal sample pot (Global Scientific).
2. Remaining heifers had faecal samples collected immediately after defecation; these heifers were monitored by a researcher in their normal environment. When these heifers were observed to defecate, a freshly voided sample was collected into a 30 ml universal container with a spoon (Global Scientific) with extreme care not to contaminate the sample (i.e. a small amount of faeces was collected from the top of the faecal pat).

### Sample processing

Samples were refrigerated from collection to processing. Once in the lab, samples were transferred from the collection device into individual labelled sterile stomacher bags and suspended in 10 mL/g of phosphate buffered saline (PBS Dulbecco A; Oxoid, Basingstoke, UK). Samples were then mixed for 1 min in a stomacher (Stomacher 400, Seward, Worthing, UK). Samples were mixed 50:50 v/v with 100% sterile glycerol and aliquots stored at -80°C until further processing. All samples were processed and frozen within 5 days of collection. Half of the samples were then defrosted for microbiology processing within 6 months of collection. The other half were stored frozen for a further 6 months (up to 12 months total) before defrosting and microbiological analysis.

### Microbiology

Each sample was defrosted at room temperature and 0.5 mL of the thawed sample was added to 9.5 mL lauryl sulphate broth. This mixture was then incubated at 37°C for 16-24 h.

Broth-enriched samples were plated – both without dilution and with 1:100 dilution - onto TBX agar plates containing 16 mg.L^-1^ tetracycline, 8 mg.L^-1^amoxicillin or 16 mg.L^-1^ cefalexin, meaning that growth indicated bacteria resistant to the European Committee on Antimicrobial Susceptibility Testing breakpoint concentration, as used in the OH-STAR study (Schubert *et al*., 2021). Plates were then incubated at 37°C for 16-24 h. Plates were assessed for the presence or absence of blue (*E. coli*) individual colonies, creating binary results. If there was overgrowth even at 1:100 dilution, serial dilutions were plated out to check for the presence of individual colonies.

### Data analysis

All data analysis was performed using R (https://www.r-project.org/). Our objective was to test associations between sample-level positivity for *E. coli* resistant to each of 3 test antibacterials and the type of individual heifer samples (rectal or pat), the sample storage time (<6 months or 6-12 months), sample-level positivity for resistant

*E. coli* in environmental samples collected from around the test heifers in the 12 months prior to individual heifer sampling, as defined in the OH-STAR study (Schubert *et al*., 2021), as well as farm ABU (as measured in the OH-STAR study).

The dependent variable was the presence or absence of *E. coli* resistant, separately, to each of the 3 test antibacterials in each individual heifer faecal sample. Analysis was performed using multilevel, multivariable logistic regression with farm as a random effect and had the following fixed effects:

- Whether the sample was from a rectal examination or a freshly voided faecal pat
- Time of post-sample processing (retrieval from the freezer and microbiological analysis); within 6 months *vs*. 6-12 months of storage
- Total ABU on the farm for the first 12 months of the project after the farm was enrolled (a 12-month period between January 2017 – May 2018 depending on the date of enrolment for different farms), as measured in mg/ PCU (ESVAC, 2015)
- Proportion of environmental samples which were positive for resistance to that antibacterial; environmental samples were collected monthly on each farm for the 12 months preceding heifer sampling as part of the OH-STAR study
- Additional ABU variables for specific antibacterials, depending on the dependent variable (using prior knowledge of what antibacterial agents could select for resistance to the outcome variable (Schubert *et al*., 2021)). Specifically, for the amoxicillin model, the amount of penicillins and potentiated amoxicillins (measured in mg/PCU) used on the farm were modelled as well as total amoxicillin use; for cefalexin, the amount of 3^rd^- and 4^th^-generation as well as 1^st –^ generation cephalosporin use (in mg/PCU) was also modelled. The total amount of tetracycline used on the farm was modelled in the tetracycline model.

Within the models, continuous variables were centred and scaled using the scale function in R so that they had a standard deviation of 1. Models were initially run with all fixed effects included and then further refined using a backwards stepwise procedure (Dohoo *et al*., 2010) until only variables with p<0.05 remained in the model. Variables which were retained in the model after the backwards stepwise procedure were checked for multicollinearity by removing each variable in turn and checking that the confidence intervals for the estimate for each variable still overlapped. Predictive accuracy was checked using area under the Receiver Operating Characteristic Curve; good predictive accuracy was evident for all models (amoxicillin model - 0.86; tetracycline model - 0.89; cefalexin model - 0.79).

## Results

One sample was collected from each of 364 individual heifers housed on 37 dairy farms. Samples were either collected directly from the rectum or were from freshly voided faecal pats accurately linked to an individual animal through observation by a researcher. The proportions of samples positive for *E. coli* resistant to either of the 3 test antibacterials (cefalexin, amoxicillin or tetracycline) at the farm as well as at the sample level are shown in **Table 1**. Three of the 37 farms had no resistance to any of the 3 test antibacterials detected in any of the individual heifer samples collected; one of the 37 farms had resistance detected to all 3 antibacterials in all individual heifer faecal samples.

**Table 1.** Number (N) and percentage of individual heifer samples (one sample collected per heifer) showing resistance to amoxicillin, cefalexin and tetracycline at the farm and sample levels

| Antimicrobial | Resistance: Farm level |  | Resistance: Sample level |  |
| --- | --- | --- | --- | --- |
|  | N | % | N | % |
| Amoxicillin | 29/37 | 78 | 114/364 | 31 |
| Tetracycline | 25/37 | 68 | 96/364 | 26 |
| Cefalexin | 29/37 | 78 | 82/364 | 23 |

The results of models predicting sample-level resistant *E. coli* positivity (**Tables 2-4**) showed that for each of the 3 test antibacterials, individual heifer samples collected directly from the rectum were more likely to be positive for resistant *E. coli* than those collected from freshly voided faecal pats. The weakest effect was for cefalexin resistance (**Table 4**) and the strongest for tetracycline resistance (**Table 3**), where the odds of rectal samples being positive for resistant *E. coli* were >8 times greater than faecal pats. Storage of the processed samples for 6-12 months in a freezer at - 80°C was associated with significantly reduced sample-level positivity for amoxicillin (**Table 2**) or tetracycline (**Table 3**) resistant *E. coli* compared to those stored for <6 months. There was no difference for cefalexin resistance with regards to storage.

**Table 2.** Risk factors for the presence of amoxicillin resistance in individual heifer faecal samples taken from 37 farms.

| Risk factor | Odds ratio | 95% confidence interval |  | p-value |
| --- | --- | --- | --- | --- |
|  |  | lower | upper |  |
| Rectal sample | 3.82 | 1.67 | 8.77 | 0.002 |
| Freshly voided sample | reference |  |  |  |
| 6-12 months storage | 0.28 | 0.11 | 0.66 | 0.004 |
| <6 months storage | reference |  |  |  |

**Table 3.** Risk factors for the presence of tetracycline resistance in individual heifer faecal samples taken from 37 farms.

| Risk factor | Odds ratio | 95% confidence interval |  | p-value |
| --- | --- | --- | --- | --- |
|  |  | lower | upper |  |
| Rectal sample | 8.17 | 3.47 | 19.21 | <0.000001 |
| Freshly voided sample | reference |  |  |  |
| 6-12 months storage | 0.18 | 0.07 | 0.46 | 0.0003 |
| <6 months storage | reference |  |  |  |
| Total farm ABU^ in the first 12 months of the project measured in mg/PCU* | 2.09 | 1.18 | 3.71 | 0.01 |

^ABU = antibacterial use; \*PCU = population correction unit as used for the standard UK AMU reporting metric (ESVAC, 2015)

**Table 4.** Risk factors for the presence of cefalexin resistance in individual heifer faecal samples taken from 37 farms.

| Risk factor | Odds ratio | 95% confidence interval |  | p-value |
| --- | --- | --- | --- | --- |
|  |  | Lower | Upper |  |
| Rectal sample | 2.05 | 1.04 | 4.04 | 0.04 |
| Freshly voided sample | reference |  |  |  |

Positivity for *E. coli* resistant to a particular antibacterial in environmental samples collected close to the 364 test heifers monthly over the 12 months prior to individual heifer sample collection had no significant association with sample-level positivity for the same antibacterial in the individual heifer samples. There was no significant association between the use of tetracycline, cefalexin or amoxicillin and sample level positivity for resistance to either of these drugs. A significant association with an ABU metric was only seen for tetracycline resistance. After accounting for the fact that numerical values were centred and scaled, the tetracycline model showed a positive association between the odds of an individual heifer sample being positive for tetracycline-resistant *E. coli* and total ABU on the farm in the 12 months prior to heifer sample collection (OR: 2.09). Specifically, this means that the odds of a sample being positive for tetracycline-resistant *E. coli* increased by 2.09 for every 37.6 mg/PCU increase in total ABU on a farm (**Table 3**).

We were concerned to note the reduced odds of finding resistant *E. coli* in samples stored for 6-12 months, so we re-analysed environmental samples from our previously published OH-STAR project (5, 20) which had been processed and stored at -80°C in 50% glycerol exactly as were the samples in this parallel individual heifer sampling study. We chose 20 environmental samples collected in January-March 2017 that had been positive with high numbers (>1000 cfu/mL) of cefotaxime-resistant *E. coli* and which were known (based on multiplex PCR) to have the pMOO-32 plasmid, carrying *bla*_CTX-M-32_, conferring cefotaxime resistance (Findlay *et al*., 2021). Samples were recovered from the freezer in November 2019. Nine (45%) of the samples were positive for cefotaxime-resistant *E. coli* but only one of these samples carried cefotaxime-resistant *E. coli* at a density of >200 cfu/mL. Based on the pMOO-32-specific multiplex PCR, only this single sample with the highest density grew cefotaxime-resistant *E. coli* carrying pMOO-32.

## Discussion

The overall positivity rate for resistant *E. coli* being found in individual heifer samples in this study was lower at the farm level (78, 68, 78% farms with at least one positive sample for amoxicillin, tetracycline and cefalexin resistant *E. coli*, respectively) than that seen in our parallel OH-STAR study (Schubert *et al*., 2021) of environmental samples (100% farms with at least one sample positive for amoxicillin, tetracycline and cefalexin resistant *E. coli*, respectively). The percentage of samples positive for resistant *E. coli* across the whole study (31, 26 and 23% for amoxicillin, tetracycline and cefalexin, respectively) was also lower than in the OH-STAR study (Schubert *et al*., 2021) for amoxicillin (66%) and tetracycline (69%) but was very similar for cefalexin (19%). Many studies, including OH-STAR, have found that areas of a farm housing calves and youngstock generally are associated with higher prevalence of ABR (Hancock *et al*., 1997, Horton *et al*., 2016, Schubert *et al*., 2021). The findings of this study suggest that 18-month-old heifers may have resistance profiles more comparable with the adult herd than with pre-weaned calves, which may explain why sample-level positivity rates were lower in this study than in the OH-STAR study, which included environmental samples associated with cattle of all ages (Schubert *et al*., 2021). Another explanation for this observed lower prevalence of resistance (despite the use of enrichment culture here, which should increase the chances of finding resistant bacteria) is that, in OH-STAR, environmental samples were likely representative of faeces from multiple animals, meaning the chances of finding resistance might have been higher. The implication here is that the sample-level prevalence of resistance relates to the number of animals excreting resistant bacteria and the abundance of resistant bacteria excreted by the positive animal(s).

This study found, in all models, greater odds of isolating resistant *E. coli* from samples that were collected directly from the rectum of heifers compared to samples collected from freshly voided faecal pats. An initial assumption would be that the freshly voided faecal pat samples may have been subject to exposure to temperatures below body temperature, resulting in a lower prevalence of resistance in these samples, since reduced temperature has been associated with reduced odds of finding resistant *E. coli* by ourselves and other authors (MacFadden *et al*., 2018, Schubert *et al*., 2021). However, the fact that animals were observed defecating and the faecal pat sample was then collected almost immediately make this possibility seem unlikely, and the explanation may simply be the relationship with the position within the rectum where the bacteria reside immediately prior to sampling. For example, rectal examination may dislodge bacteria adhered to the surface of the rectum which are not normally excreted. This hypothesis will require further analysis, but the fact that certain *E. coli* reside in different parts of the gastrointestinal tract has been shown previously (Grauke *et al*., 2002, Naylor *et al*., 2003, Low *et al*., 2005), as has the fact that recto-anal mucosal swab sampling has a different sensitivity to faecal cultures for determining *E. coli* O157 prevalence in cattle (Greenquist *et al*., 2005). One other study has compared the prevalence of resistant *E. coli* in rectal versus pat samples, and these authors did not find a difference for tetracycline resistance (along with other antibacterials non-comparable to this study (Wagner *et al*., 2002). Importantly, however, faecal pat samples in that study were collected from the pen floor of feedlot cattle (Wagner *et al*., 2002). It is possible, therefore, that the faecal pat samples more closely represented the environmental samples from our OH-STAR study, which had a higher prevalence of resistance (Schubert *et al*., 2021), likely because they were not actually individual samples but aggregates of multiple positive and negative faecal pats.

In this study, the storage of samples for 6-12 months (in comparison to <6 months) at -80°C was associated with lower odds of finding *E. coli* resistant to amoxicillin and tetracycline. Furthermore, re-analysis of OH-STAR environmental samples stored for 2.5 years at -80°C and known to be strongly positive for *E. coli* carrying the pMOO-32 plasmid dominant as a cause of 3^rd^-generation cephalosporin resistance in *E. coli* in these study farms revealed that only 1/20 were still positive for this plasmid after storage. The presence of ABR in *E. coli* has been associated with a fitness cost to the bacteria (Basra *et al*., 2018) and warmer temperatures have been suggested to result in increased growth rates of bacteria with ABR phenotypes (MacFadden *et al*., 2018). Storage at -80°C, as in this study, is intended to stop all growth of bacteria and so remove competition. However, in this case, resistant bacteria were not recoverable after such storage of 19/20 samples. Research needs to be done, therefore, to identify conditions/media that allow long-term storage of cattle faecal samples, important for follow-up surveillance projects and comparisons between samples collected in longitudinal surveillance studies, without loss of ABR. We would urge caution for others carrying out related environmental studies of ABR prevalence and hoping to store samples in the long term.

This study also showed an association between total ABU on the farm (as measured in mg/PCU) with increased odds of finding tetracycline-resistant *E. coli* in the individual heifer faecal samples. Total ABU was not found to be associated with sample-level positivity for amoxicillin- or cefalexin-resistant *E. coli*. There was no association found with other ABU variables that were included in the models, including usage of cefalexin, amoxicillin and tetracycline, separately, as potential drivers of resistance to the relevant antibacterial. Importantly, the OH-STAR study of environmental samples on these farms also found that only the odds of finding samples positive for tetracycline-resistant *E. coli* were associated with total ABU on farms; no significant associations were found between ABU and resistance to amoxicillin (cefalexin resistance was not modelled in that study) (Schubert *et al*., 2021). An increased odds (OR 2.09) of finding tetracycline-resistant *E. coli* in the individual heifer samples in this study associated with a farm-level increase of 37.6 mg/PCU must be considered in the context that the average total ABU on UK dairy farms has been calculated to be 22.5 mg/PCU in 2019 in the Veterinary Antibiotic Resistance Sales and Surveillance report (VARSS, 2019), so our findings are unlikely to be predictive of ABR detection changes associated with smaller changes in ABU within a single farm. One study, however, found ABU on UK dairy farms to range from 0.36 – 97.7 mg/PCU (Hyde *et al*., 2017). Therefore, the risk of finding tetracycline-resistant *E. coli* in faecal samples from heifers could feasibly be greater on UK dairies with the highest ABU relative to those with the lowest ABU.

Antibacterial use in dairy cows has been shown to be largely driven by mastitis treatments and dry cow therapies (Stevens *et al*., 2016, Hyde *et al*., 2017) however, the use of parenteral treatments for mastitis have been associated with the presence of ABR *E. coli* in dairy herds (Santman-Berends *et al*., 2017). Treatments for intramammary infection (and other production diseases) would not commonly be necessary for non-lactating heifers, so it may be that this is one reason why ABU was not associated more widely with the odds of finding ABR *E. coli* in the individual heifer samples in this study. At a national level, higher total ABU has been found to be associated with increased prevalence of resistance to certain antibacterials found in livestock-associated *E. coli* (Dorado-Garcia *et al*., 2016, Callens *et al*., 2018). However, most studies that assess the association between total farm ABU and ABR patterns suggest that total farm ABU is generally a poor predictor of specific resistance patterns on individual farms (Berge *et al*., 2010, Gonggrijp *et al*., 2016, Santman-Berends *et al*., 2017, Gay *et al*., 2019). It is likely that multiple factors affect the prevalence of ABR on a farm and that the interactions of these variables are revealed in different ways by different studies.

No association was seen between the odds of an individual heifer sample being positive for *E. coli* resistant to an antibacterial, and the sample-level prevalence of *E. coli* resistant to that same antibacterial in environmental samples collected monthly in the previous 12 months from the farm where the individual heifer sample was also collected. This result would make it seem that heifers are not consistently influenced by the microbiology of the environment they inhabit. Environmental sampling using over-boot socks has been shown to correlate with individual animal sampling performed at the same time in one study (Agren *et al*., 2018), however that study only analysed herd-level *Salmonella* detection rather than quantitatively analysing ABR as we did in the present study. The prevalence of ABR detected in the environmental samples from the farms under study here may have been influenced by the limit of detection as enrichment culture was not used; the prevalence of ABR detected in the environmental samples would also have been representative of an aggregate of many animals. Nonetheless, this reinforces the suggestion - based on individual heifer samples with a very low limit of detection (approximately 20 cfu/g of faeces) - that ABR prevalence in a group of heifers is driven by the number of animals excreting resistant bacteria rather than the density of resistant bacteria excreted by each member of the herd.

Overall, this study demonstrates that sampling methodology and sample handling had more consistent and more significant associations with ABR *E. coli* detection in individual animal faecal samples from dairy heifers than management and husbandry factors, including ABU on the farm. Most importantly, this study has highlighted that when investigating ABR prevalence on farms or in individual animals, using consistent methodology within and between studies is crucial. Factors that have been identified in other studies such as temperature, location on the farm and whether the sampling was done indoors or out at pasture are also important to factor in (Schubert *et al*., 2021). Further consideration should also be given to whether samples were collected directly from the rectum or from faecal pats, whether these are truly individual samples (if that is desired), and how the samples were processed or stored prior to ABR testing.

## Acknowledgements

We want to thank all the farmers who participated in this study. In addition, we would like to thank the milk processing companies who supported the recruitment process for the OH-STAR study. This work was funded by grant NE/N01961X/1 to M.B.A & K.K.R from the Antimicrobial Resistance Cross Council Initiative supported by the seven United Kingdom research councils. J.E.S. is supported by a scholarship from the Medical Research Foundation National PhD Training Programme in Antimicrobial Resistance Research (MRF-145-0004-TPG-AVISO).

## Transparency declarations

The authors declare no conflicts of interests. Farming and veterinary businesses who contributed data and permitted access for sample collection were not involved in the design of this study or in data analysis and were not involved in drafting the manuscript for publication.

## Author Contributions

Conceived the Study: K.K.R., M.B.A.

Collection of Data: H.S., E.F.P., J.E.S., supervised by T.A.C., K.K.R., M.B.A.

Cleaning and Analysis of Data: A.T., H.S., E.F.P., J.E.S. V.C.G., supervised by K.K.R., M.B.A.

Initial Drafting of Manuscript: A.T., H.S., J.S., M.B.A

Corrected and Approved Manuscript: All authors

